# A spinal synergy of excitatory and inhibitory neurons coordinates ipsilateral body movements

**DOI:** 10.1101/2023.03.21.533603

**Authors:** Marito Hayashi, Miriam Gullo, Gokhan Senturk, Stefania Di Costanzo, Shinji C. Nagasaki, Ryoichiro Kageyama, Itaru Imayoshi, Martyn Goulding, Samuel L. Pfaff, Graziana Gatto

**Author notes:** Co-corresponding authors Contacts: Itaru Imayoshi, Institute for Frontier Life and Medical Sciences, Martyn Goulding, Salk Institute for Biological Studies, Sam Pfaff, Salk Institute for Biological Studies, Graziana Gatto, University Hospital of Cologne.

## Abstract

Innate and goal-directed movements require a high-degree of trunk and appendicular muscle coordination to preserve body stability while ensuring the correct execution of the motor action. The spinal neural circuits underlying motor execution and postural stability are finely modulated by propriospinal, sensory and descending feedback, yet how distinct spinal neuron populations cooperate to control body stability and limb coordination remains unclear. Here, we identified a spinal microcircuit composed of V2 lineage-derived excitatory (V2a) and inhibitory (V2b) neurons that together coordinate ipsilateral body movements during locomotion. Inactivation of the entire V2 neuron lineage does not impair intralimb coordination but destabilizes body balance and ipsilateral limb coupling, causing mice to adopt a compensatory festinating gait and be unable to execute skilled locomotor tasks. Taken together our data suggest that during locomotion the excitatory V2a and inhibitory V2b neurons act antagonistically to control intralimb coordination, and synergistically to coordinate forelimb and hindlimb movements. Thus, we suggest a new circuit architecture, by which neurons with distinct neurotransmitter identities employ a dual-mode of operation, exerting either synergistic or opposing functions to control different facets of the same motor behavior.

## Introduction

Walking in terrestrial vertebrates is generated by the hierarchical interaction of supraspinal, sensory and spinal circuits that stabilizes posture and ensures the fluid execution of locomotion (Grillner and El Manira, 2020). The dynamic interplay among these three systems preserves balance as the body locomotes forward, via the active coordination of trunk and appendicular muscles across the whole body. Thus, the nervous system not only encodes the dynamics of individual joints, but also harmonizes motion across joints and limbs to avoid postural instability. Successful coordination of all four limbs during locomotion needs to flexibly account for multiple parameters of limb movement in order to maintain body stability: a) the forward distance covered by each step (stride), b) the stride frequency (cadence), c) the time spent supporting the body (stance), and d) the time spent propelling it forward (swing). Limb movements are coordinated across left and right sides of the body (bilateral coupling) and between forelimbs and hindlimbs (ipsilateral and diagonal coupling) **(Figure 1A)**. Interlimb coupling is flexibly phased during locomotion to accommodate changes in gait (Bellardita and Kiehn, 2015), with for example bilateral limbs transitioning from alternation during trot to synchrony during bound **(Figure 1A)** (Bellardita and Kiehn, 2015; English and Lennard, 1982; Grillner and El Manira, 2020; Lemieux et al., 2016). These interlimb patterns are evolutionary conserved mechanisms observed also in bipedal species. Synchronized arm and leg movements in humans facilitate faster running paces and stabilize the center of gravity (Catavitello et al., 2018; Dietz, 2002). Nevertheless, the neuronal substrates coordinating movement between forelimbs and hindlimbs remain largely unknown.

**Figure 1.**
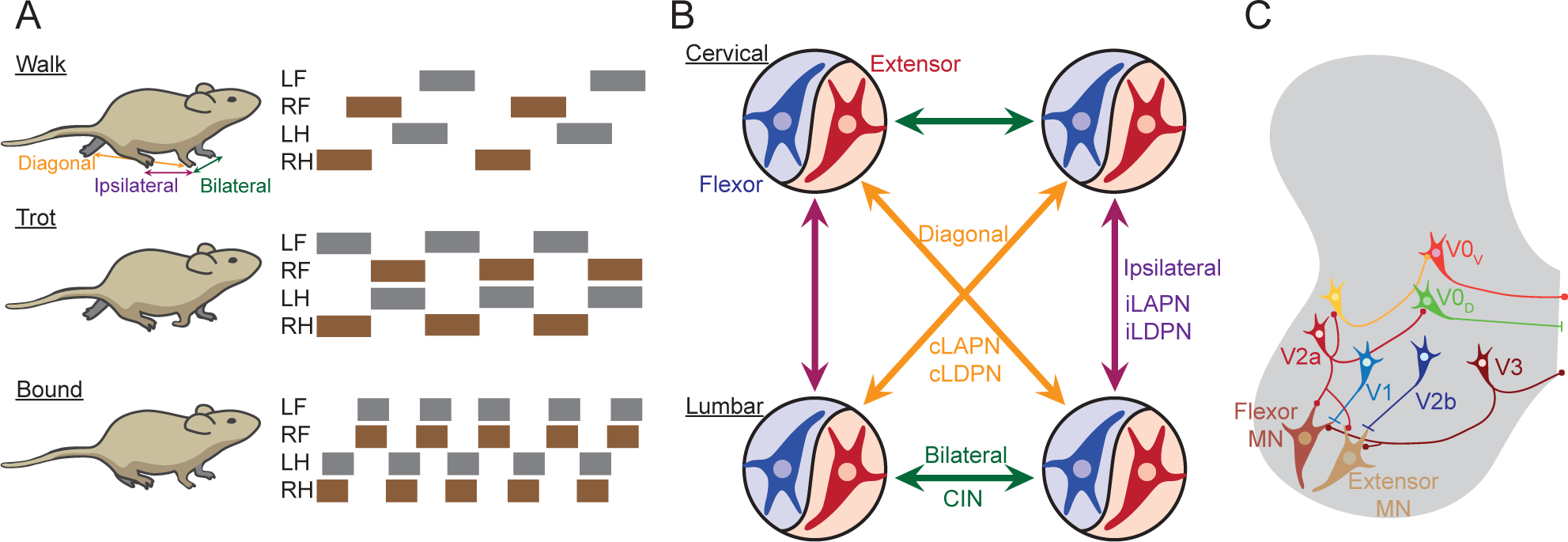
Body coordination during locomotion. (A) Schematic illustrating three different gaits in mice. The colored rectangle represents the time each limb is in contact with the ground. Each gait is characterized by different coupling of left forelimb (LF), right forelimb (RF), left hindlimb (LH) and right hindlimb (RH). (B) Schematic illustrating the anatomical connections across half-center modules. Each module is composed of a pool of excitatory and inhibitory neurons controlling extensor (red) and flexor (blue) muscles. Commissural pathways (CIN) at each girdle (green arrows) ensure left and right coordination, ipsilateral (violet) ascending (iLAPN) and descending (iLDPN) propriospinal pathways contribute to the unilateral coordination of forelimb and hindlimb, whereas commissural (orange) ascending (cLAPN) and descending (cLDPN) propriospinal tract connect opposite forelimb and hindlimb. (C) Schematic illustrating the main classes of premotor neurons forming the central pattern generator (CPG) circuits: the inhibitory ipsilateral neurons (V1 and V2b), the excitatory ipsilateral neurons (V2a), the inhibitory commissural neurons (V0D), the excitatory commissural neurons (V0V) and the dual-projecting excitatory neurons (V3). V1 and V2b neurons preferentially inhibit flexor and extensor motoneurons (MNs), respectively.

Rhythmic movements of individual limbs are controlled by dedicated premotor locomotor networks, the half-center modules (Grillner, 1975; Kiehn, 2006). Each half-center module contains the microcircuit for the coordination of flexor and extensor muscles that secures stance and swing alternation (Brown, 1911; Grillner and El Manira, 2020; Krouchev et al., 2006). Half-center modules are connected by local commissural axons, which ensure the appropriate bilateral coordination (Kiehn, 2016; Lanuza et al., 2004; Talpalar et al., 2013). Commissural and ipsilateral propriospinal projections between the cervical and lumbar spinal cord facilitate the diagonal and ipsilateral coordination of forelimbs and hindlimbs, respectively **(Figure 1B)** (Pocratsky et al., 2017; Ruder et al., 2016; Zhang et al., 2021). This modular and multi-layered organization provides a flexible infrastructure for the dynamic spatiotemporal coordination of the limbs, preventing mid-air collisions, paw trampling and postural instability.

Although the major spinal cell types composing the premotor network that regulates locomotor coordination have been identified (Goulding, 2009), the neuronal substrates and propriospinal pathways that couple and coordinate movement across all four limbs remain largely unknown. The premotor locomotor network, also known as central pattern generator (CPG), comprises six cardinal neuron populations: V1, V2a, V2b, V3, V0, dI6 neurons that are distinct in their genetic origin, connectivity and neurotransmitter expression **(Figure 1C)** (Goulding, 2009; Grillner and Jessell, 2009). The ipsilaterally projecting inhibitory V1 and V2b neurons cooperate to regulate flexion and extension alternation and the timing of stance and swing phasing (Bourane et al., 2015; Zhang et al., 2014), thus functioning as integral parts of the half-center modules. The commissural V0 neurons, the ipsilateral V2a neurons and the dual-projecting V3 neurons coordinate limb-coupling and the transition to faster gaits (Bellardita and Kiehn, 2015; Crone et al., 2008, 2009; Lanuza et al., 2004; Talpalar et al., 2013; Zhang et al., 2021). For example, when the activity of V0, V2 or V3 neurons is disrupted, mice adopt markedly altered or unstable gaits as locomotor speed increases (Bellardita and Kiehn, 2015; Crone et al., 2008, 2009; Lanuza et al., 2004; Talpalar et al., 2013; Zhang et al., 2021). Nevertheless, these disruptions do not alter interlimb coordination during self-paced locomotion, suggesting that this facet of motor control might be regulated by more complex neural dynamics involving multiple cardinal classes.

Here, we analyzed the combinatorial function of excitatory (V2a) and inhibitory (V2b) neurons that derive from the common spinal V2 progenitor domain (Alaynick et al., 2011; Goulding, 2009; Kiehn, 2016). We leveraged the transient and restricted expression of *Hes2* in V2 progenitors to generate a new genetic tool (*Hes2^iCre^*) and assess the contribution that the spinal V2 lineage neurons make to intra-and interlimb coordination during locomotion. Developmental silencing of the Hes2 neurons in mice dramatically changed the walking gait without affecting intralimb coordination. Inactivation of Hes2 neurons caused an adaptive gait, with increased cadence and reduced stride length, to compensate the postural instability induced by the malfunction of these neurons. This postural instability is illustrated by the increased time that Hes2 neuron-silenced mice spent with all four limbs in stance phase during stepping and the inability of the ipsilateral hindlimb to execute forward targeting movements during swing. Furthermore, when executing more challenging motor tasks, such as narrow beam and uneven ladder, mice with inactivation of Hes2 neurons were more prone to slips, reinforcing a key role for these neurons in ensuring body balance. Using an intersectional genetic approach, we further demonstrated that these altered coordination dynamics were primarily due to the disrupted function of V2a and V2b neurons within the spinal cord. In summary, our analyses showed that the simultaneous inactivation of V2a and V2b neurons rescued the hindlimb hyper-extension observed in V2b neuron-ablated mice (Britz et al., 2015), suggesting that these two neuron populations act antagonistically to pattern flexor and extensor alternation. Intriguingly, our findings also revealed that spinal excitatory V2a and inhibitory V2b neurons act synergistically to modulate interlimb spatiotemporal dynamics, forming a specialized hub for ipsilateral body coordination. Thus, we identified a microcircuit of spinal excitatory and inhibitory neurons that by functioning in antagonism and synergy can control distinct facets of locomotor coordination.

## Results

### Identification of a novel marker of the V2 neuron lineage

The embryonic V2 spinal progenitor domain differentiates into excitatory V2a and inhibitory V2b neurons (Alaynick et al., 2011; Goulding, 2009; Kiehn, 2016), via the Delta-Notch signaling pathway (Del Barrio et al., 2007; Kimura et al., 2008; Peng et al., 2007; Yang et al., 2006). These two neuronal populations share key anatomical features, including their soma positions in the ventral gray matter and ipsilateral axonal projection patterns, with subsets of V2a and V2b neurons differentiating as propriospinal neurons (Flynn et al., 2017; Ruder et al., 2016). Ipsilaterally projecting V2 propriospinal neurons likely play key roles in coordinating motor activity between forelimbs and hindlimbs. Thus, we set out to identify a novel genetic handle to target the entire V2 neuron lineage and analyze its role during locomotion. The Hes family of transcription factors is expressed in a variety of spinal progenitor domains as they undergo cell fate specification (Kageyama and Nakanishi, 1997), often acting as downstream effector of the Delta-Notch signaling cascade (Iso et al., 2003). Therefore, we performed a targeted analysis of the clusters of developing cell types in the spinal cord identified by the Briscoe lab (Delile et al., 2019), and found *Hes2* as specifically expressed in the Lhx3-marked V2 progenitor domain (Alaynick et al., 2011) (**Figure S1A**).

*Hes2* mRNA was first detected at embryonic stage 10.5 (E10.5), peaked between E11.5 and E12.5, and was extinguished by E13.5 **(Figure 2A)**. Transient expression was confined to a cluster of ventromedial cells **(Figure 2A)**, in the region where V2 progenitor cells reside (Al-Mosawie et al., 2007; Briscoe et al., 2000; Karunaratne et al., 2002). At E11.5-E12.5 *Hes2* was detected in cells near the ventricular zone, but not in the lateral mantle zone where more mature neurons reside **(Figure 2A)**. This *Hes2+* cluster was positioned dorsal to the differentiating motoneurons that are marked by the expression of Hb9 **(Figure 2B)**, *Isl1,* and *Olig2* (**Figure S1B** and **S1C**). *Hes2* detection preceded the expression of Chx10, a marker of post-mitotic excitatory V2a neurons (Ericson et al., 1997; Hayashi et al., 2018; Peng et al., 2007) **(Figure 2B)**. In addition, *Hes2* was present in cells lacking Chx10 **(Figure 2B)**, suggesting it might be also expressed in differentiating V2b neurons. Together our observations indicate that *Hes2* is transiently and specifically expressed in the V2 progenitor domain.

**Figure 2.**
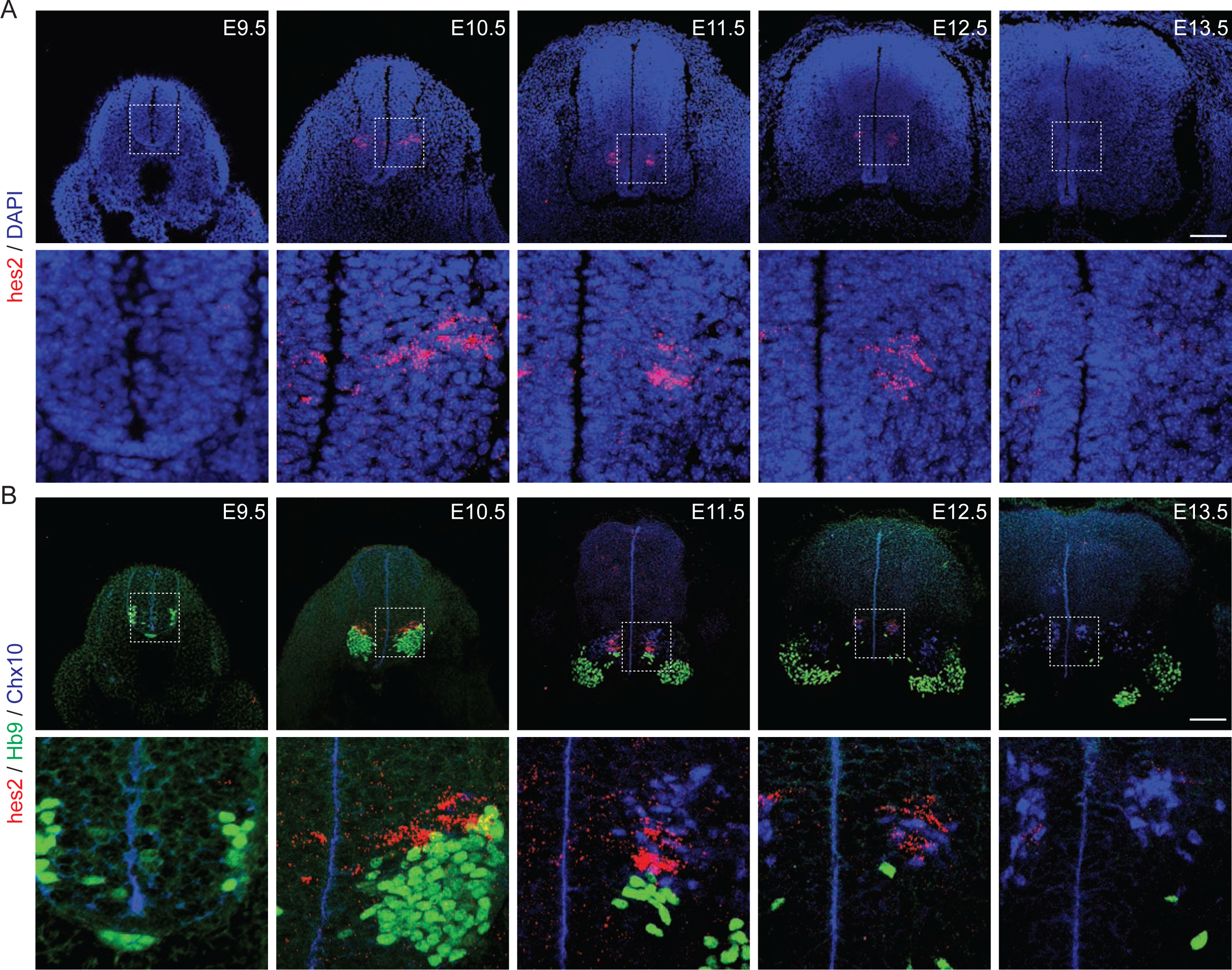
Identification of a novel marker of the spinal V2 neuron lineage. (A) Cross-sections of the developing lumbar spinal cord (E9.5 – E13.5) with labeled *Hes2* mRNA (red) and counterstained with DAPI (blue). Bottom panels are magnifications of the dotted boxes. (B) Cross-sections of the developing lumbar spinal cord (E9.5 – E13.5) stained for *Hes2* mRNA (red), Hb9 protein (green), and Chx10 protein (blue). Bottom panels are magnifications of the dotted boxes. Scale bars: 100 μm.

### Engineering a novel genetic handle to target the Hes2 neuron lineage

We generated a knock-in mouse line, in which the *iCre* recombinase is expressed under the control of the *Hes2* promoter to genetically access the entire V2 lineage (**Figure S2A-S2B**). When *Hes2^iCre^*was crossed with the *R26^LSL-LacZ^* β-galactosidase reporter, we saw that β-galactosidase signal recapitulated the expected pattern of Hes2 expression in the developing spinal cord and hindbrain at E10.5 and E11.5 **(Figure 3A)**. To better characterize the cell types labeled by *Hes2^iCre^*and confirm that it encompassed both V2a and V2b neurons, we used a fluorescent nuclear GFP reporter, *R26^LSL-Sun1-GFP^* to trace the entire Hes2 lineage, and a β-gal reporter (*Gata3^nlsLacZ^*) to mark V2b neurons **(Figure 3B)**. At E12.5 ∼48% of the GFP-expressing neurons (Hes2+) co-expressed Chx10 (V2a neurons) and ∼48% Gata3 (V2b neurons), with the remaining ∼4% negative for both markers **(Figure 3C)**. This small population of Hes2 lineage cells lacking expression of Chx10 and Gata3 were found near the ventricular zone, suggesting they are likely to be immature progenitor or precursor cells that are yet to differentiate. Almost all Chx10+ (98%) and Gata3+ (95%) neurons were targeted by the *Hes2^iCre^* (**Figure 3D** and **3E**). The missing 5% of Gata3+ neurons are the late born PKDL1+ neurons that surround the central canal (Di Bella et al., 2019), and do not express Hes2. The equal number of V2a (Chx10+) and V2b (Gata3+) neurons captured by the *Hes2^iCre^* **(Figure 3C)** and the time course of *Hes2* expression **(Figure 2A)** confirmed that this mouse line faithfully and specifically captures the entire spinal V2 neuron lineage. As *Hes2^iCre^* also captured some hindbrain cells at E11.5 **(Figure 3A)**, we further characterized its expression in supraspinal structures, using a tdTomato fluorescent reporter line (*R26^Ai14-LSL-tdTomato^*). In sections from juvenile brains we observed neurons labeled in the PAG, the pons, the superior olivary complex, and the medullary reticular nucleus (**Figure S2C**). No labeled cells were, however, detected in motor control-associated areas, such as the mesencephalic locomotor region (MLR), the motor cortex and the striatum.

**Figure 3.**
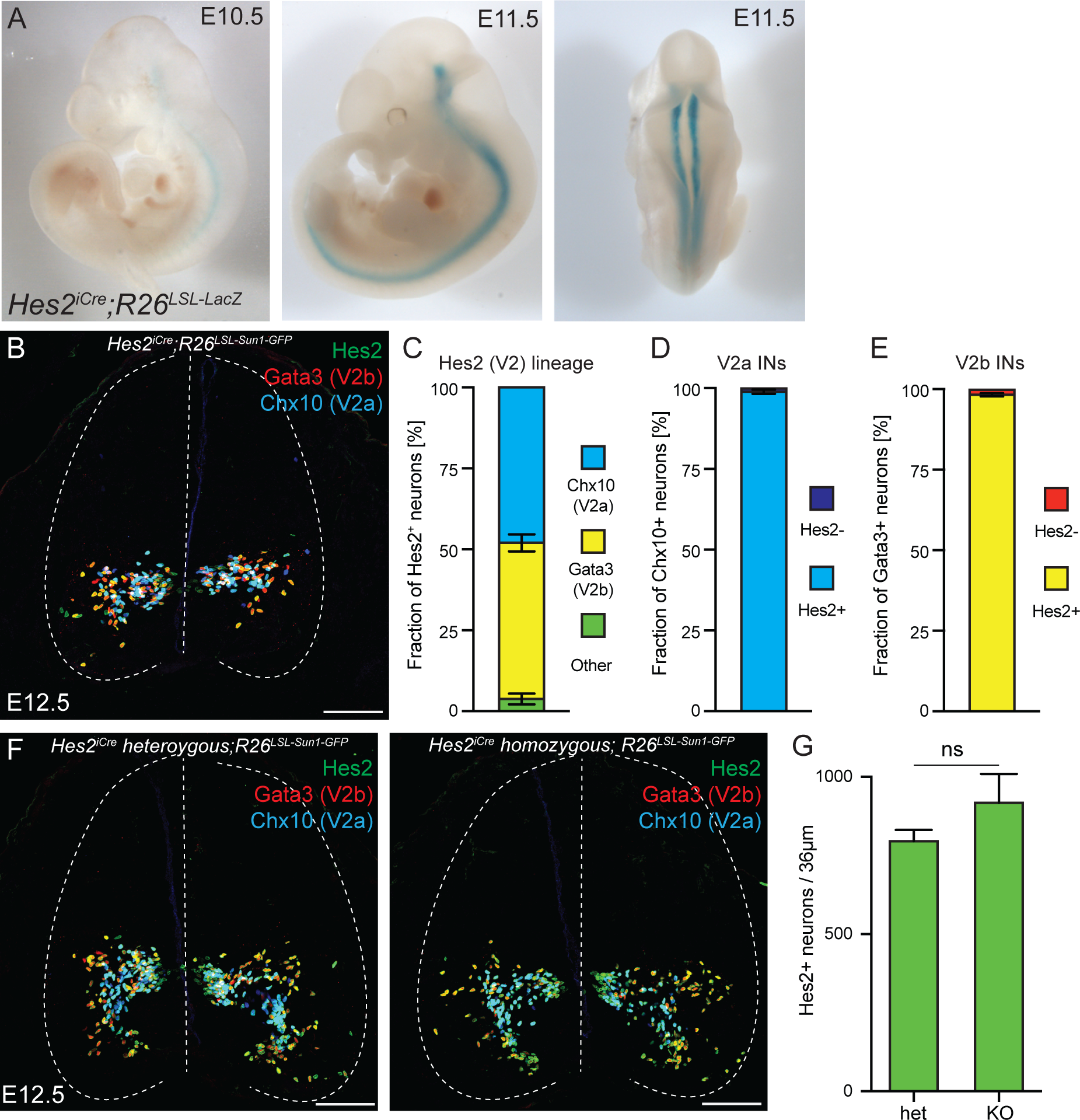
Establishing genetic access to the Hes2 neurons. (A) Whole-mount β-galactosidase staining on E10.5 and E11.5 *Hes2^iCre^;R26^LSL-LacZ^* embryos to characterize the expression of *Hes2^iCre^*. (B) Cross-sections of E12.5 lumbar spinal cord from *Hes2^iCre^;Gata3^nlsLacZ^;R26^LSL-Sun1-GFP^* embryos stained with antibodies for GFP to label Hes2 neurons (green), β-gal (*Gata3^nlsLacZ^*) to label V2b neurons (red), and Chx10 to label V2a neurons (blue). Scale bar: 100 μm. (C) Bar graph showing the percentages of V2a (Chx10+) and V2b (*Gata3^nlsLacZ^* +) neurons within the Hes2 lineage (Hes2+). (D,E) Bar graphs showing the percentage of V2a (Chx10+) (D) and V2b (*Gata3^nlsLacZ^*+) (E) neurons captured by the *Hes2^iCre^*. (F) Cross-sections of E12.5 lumbar spinal cord from *Hes2^iCre^;Gata3^nlsLacZ^;R26^LSL-Sun1-GFP^* embryos either heterozygous (left) or homozygous (right) for the *iCre* allele. The sections were stained with antibodies for GFP (green), β-gal (*Gata3^nlsLacZ^*) (red) and Chx10 (blue). Scale bars: 100 μm. (G) Bar graph showing the absence of changes in the number of Hes2 neurons in *Hes2* knockout embryos compared to heterozygous littermates, assessed by two-tailed Student’s t-test.

Next, we verified that the insertion of the *iCre* allele in the *Hes2* locus, which eliminates the Hes2 DNA binding domain (**Figure S2A**), did not disrupt spinal cord development. Heterozygous and homozygous (*Hes2* knockout) mice were generated in Mendelian ratio (**Figure S2D**), and *Hes2* knockout mice were fertile and did not display obvious behavioral deficits (data not shown). We also did not detect any difference in the numbers of Hes2 lineage (GFP+), V2a (Chx10+), or V2b (Gata3+) neurons between Hes2 homozygous and heterozygous animals (**Figure 3F**–**3G** and **S2E-S2F**). In parallel, we performed Hes2 gain-of-function experiments in the developing chick spinal cord via *in ovo* electroporation, but detected no ectopic Chx10+ (V2a) nor Gata3+ (V2b) neurons (**Figure S2G**). Thus, Hes2 function appears to be dispensable for V2 neuron development and specification despite its exquisite expression pattern, making *Hes2^iCre^* a valuable tool to study V2 neuron lineage function in adult mice.

### Developmental silencing of Hes2 neurons induces festination

Forward locomotion requires four layers of coordination: a) intralimb flexor/extensor alternation, b) intralimb coordination of proximal/distal muscles, c) left and right alternation, and d) forelimb and hindlimb coupling. In prior studies, where either excitatory V2a neurons or inhibitory V2b neurons were selectively inactivated, each population was observed to regulate a distinct facet of limb coordination (Britz et al., 2015; Crone et al., 2008, 2009; Zhang et al., 2014). V2a neurons control forelimb reaching and bilateral limb coordination during speed-dependent gait transitions (Azim et al., 2014; Crone et al., 2009), while V2b neurons contribute to regulate hindlimb flexor and extensor alternation (Britz et al., 2015), in concert with V1 neurons (Zhang et al., 2014). However, given their common developmental provenance, it is not clear whether V2a and V2b populations contribute to different functions as indicated by the individual inactivation analyses or if they also act collectively to regulate other aspects of motor coordination.

To examine the collective functions of V2a and V2b neurons, we developmentally silenced the synaptic output of Hes2 neurons by crossing *Hes2^iCre^* mice with the Cre-dependent Tetanus Toxin (TeNT) effector, *R26^LSL-TeNT^* (Zhang et al., 2008). This manipulation silenced the entire spinal V2 lineage as well as the supraspinal neurons targeted by the *Hes2^iCre^*. Littermates lacking the *Cre*allele were used as controls. Mice with inactivation of the Hes2 neurons displayed normal performance on the rotarod (**Figure S3A** and **S3B**), and normal grip strength (**Figure S3C**). However, these mice adopted an altered locomotor gait **(Figure 4A, Videos S1** and **S2**), characterized by an increased cadence of shorter strides (**Figure 4C–4F**). Both hindlimbs and forelimbs in Hes2 neuron-silenced mice displayed a shorter step cycle, in which both stance and swing phases were reduced compared to littermate controls (**Figure 4G–4K** and **S3E-S3I**). These changes in the walking pattern closely resemble the festinating gait observed in Parkinson patients (Giladi et al., 2001), characterized by the tendency to increase the frequency of stepping during locomotion (Yang et al., 2008).

**Figure 4.**
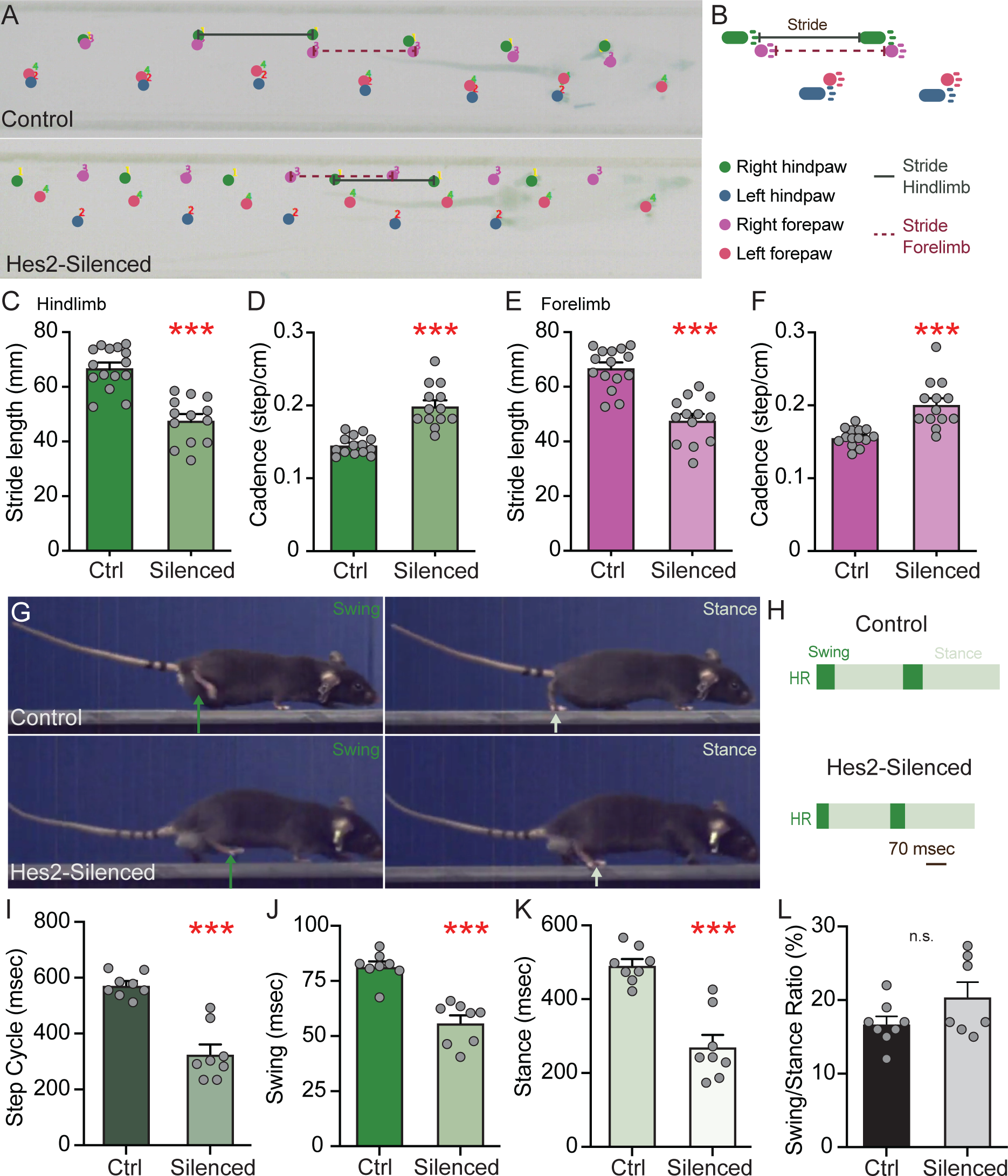
Festinating gait induced by the developmental silencing of the Hes2 neurons. (A) Bottom view of self-paced walking of mice on a runway. Paws are tracked in the indicated colors in control (upper panel) and Hes2 neuron-silenced (bottom panel) mice. (B) Schematic displaying the calculation of hindlimb and forelimb stride and the color-coding of individual limbs. (C,E) Bar graphs showing the significant shortening of the stride length of hindlimb (C) and forelimb (E) in Hes2 neuron-silenced mice compared to controls. (D,F) Bar graphs showing the significant increased stepping cadence of hindlimb (D) and forelimb (F) in Hes2 neuron-silenced mice compared to controls. (G) Representative frames showing the alternation of stance and swing phases of the hindlimb in control (upper panels) and Hes2 neuron-silenced (bottom panels) mice during self-paced walking on a wide runaway. Dark and light green arrows point to the hindlimb in stance and swing position, respectively. (H) Representative schematic showing the reduced duration of the step cycle, stance and swing for the right hindlimb (HR) in Hes2 neuron-silenced mice compared to controls. Scale bar is 70 msec. (I-K) Bar graphs showing the significant shortening of the step cycle (I), swing (J) and stance (K) duration of the hindlimb in Hes2 neuron-silenced mice compared to controls. (L) Bar graph showing the preserved ratio of stance and swing duration for each step cycle following the developmental silencing of the Hes2 neurons. Data are presented as mean ± SEM. Each mouse analyzed is represented with a gray filled circle. Statistical analysis was done using two-tailed Student’s *t-test*.

Notably, the stance and swing ratio within each step cycle was not altered in Hes2 neuron-silenced mice, demonstrating correct flexor and extensor coordination and intact functioning of the half-center modules **(Figure 4L)**. This was confirmed by performing additional behavioral tests and electromyographic (EMG) recordings in freely moving animals. Silencing the Hes2 neurons did not alter flexor and extensor muscle alternation, as indicated by the mutually exclusive firing patterns of tibialis anterior (flexor) and gastrocnemius (extensor) muscles during locomotion (**Figure S3J** and **S3K**), and the unchanged latency between antagonist muscle firing (**Figure S3L**). The correct execution of sensory-evoked behaviors (pinprick- and heat-induced withdrawal reflex) as sequence of flexion (lift-up the paw) and extension (placement of paw back to the ground) further confirmed our hypothesis that the Hes2 neurons are dispensable for intralimb coordination (**Figure S3M** and **S3N**).

In summary, developmental silencing of the Hes2 neurons, which include the spinal V2 neuron lineage, leads to festination. Animals with Hes2 neuron inactivation walk with increased cadence of stepping and shortened strides, but preserve flexor and extensor muscle alternation and intralimb coordination.

### Silencing Hes2 neurons modifies the spatial and temporal dynamics of interlimb coordination

The short strides and increased cadence observed in Parkinsońs patients are thought to develop as an adaptive mechanism to maintain a stable center of gravity, as the affected individuals begin to walk with their trunk involuntarily leaning forward (Giladi et al., 2001). We therefore hypothesized that the festination-like gait in mice with Hes2 neuron silencing might function to compensate for postural instability during locomotion. To further assess the causes for the shorter strides in Hes2 neuron-silenced animals, we examined in more detail the spatial **(Figure 4A)** and temporal (**Figure 5A** and **5B**) dynamics of the bilateral, diagonal and ipsilateral limb coordination. Footprint analysis during self-paced locomotion revealed that Hes2 neuron-silenced mice keep their hindlimbs closer to the body (**Figure S4B**), and their forelimbs and hindlimbs closer on the diagonal axis (**Figure S4C**). The increased distance observed between the ipsilateral forelimbs and hindlimbs following Hes2 neuron silencing (**Figure S4D**) is consistent with the shortened stride length of the individual limbs (**Figure 4C** and **4E**). In control mice, the hindlimb typically moves forward to occupy the position previously occupied by the ipsilateral forelimb, however, in Hes2 neuron-silenced mice the hindlimb was seen to land short of the forelimb spot (**Figure 4A, 5A** and **5F-5G**). Taken together, these changes in interlimb coordination are consistent with mice keeping the limbs closer to the body, limiting forward movements in the attempt to preserve body stability. Next, we examined whether the changes in limb spatial positioning were also paralleled by modifications in the temporal dynamics of interlimb coordination. We reasoned that if silencing of the Hes2 neurons caused postural instability, mice should spend more time with all four paws on the ground during locomotion. As predicted, we observed that mice with Hes2 neuron inactivation walked with frequent intervals, in which all four limbs were contemporaneously in their stance (support) phase, while littermate controls locomoted predominantly keeping only three paws on the ground (**Figure 5A–5C**). This increase in time spent on four-paw support confirms our hypothesis that Hes2 neuron-silenced mice alter their gait to optimize their postural stability.

**Figure 5.**
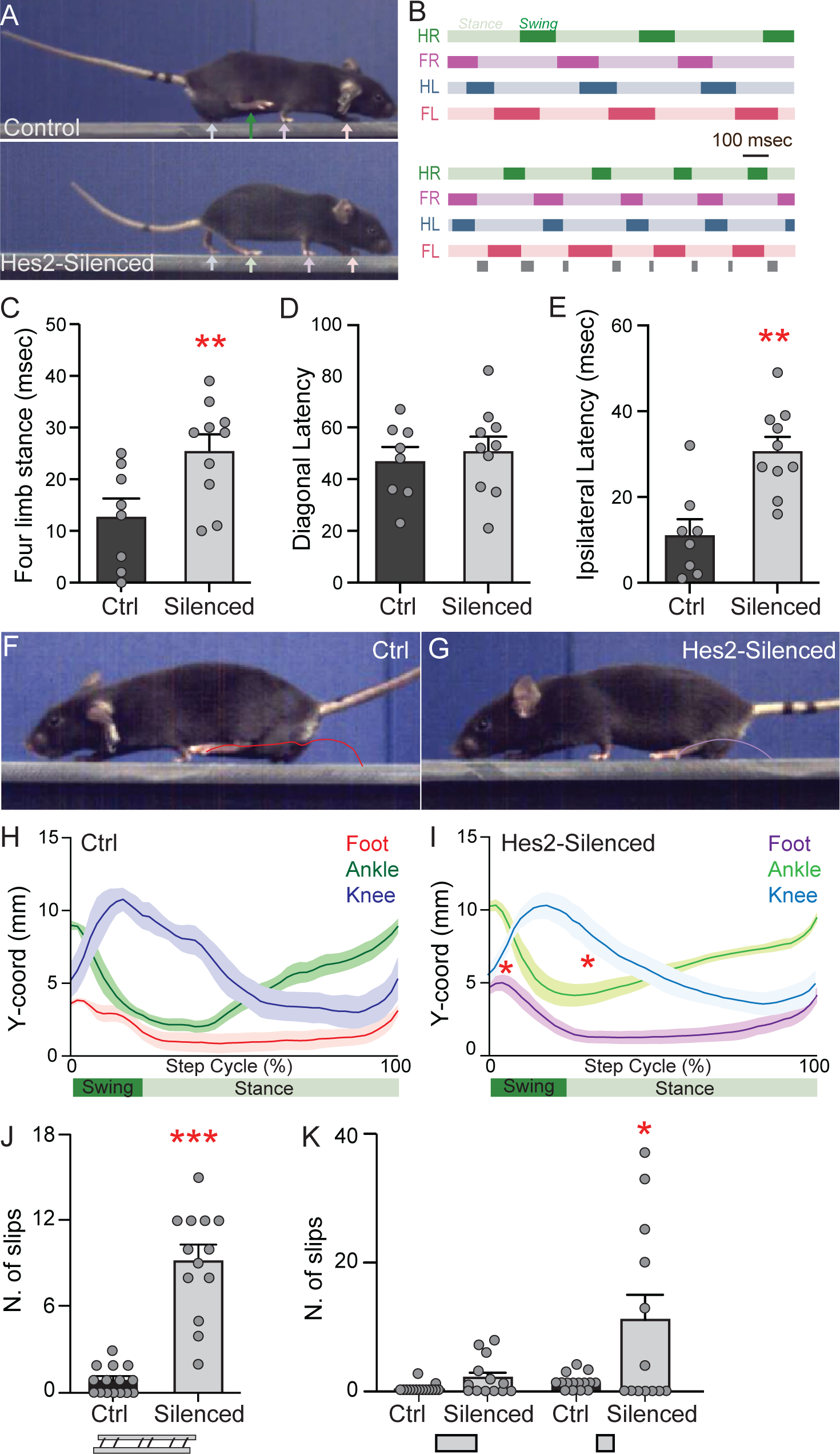
Disruption of ipsilateral body coordination by developmental silencing of the Hes2 neurons. (A) Representative frames showing the limb positioning in control (upper panel) and Hes2 neuron-silenced (bottom panel) mice during self-paced walking on a wide runaway. Dark and light green arrows point to the hindlimb in stance and swing position, respectively. (B) Representative schematics showing the temporal dynamics of interlimb coordination in control (upper panel) and Hes2 neuron-silenced (bottom panel) mice during self-paced walking. Note the extended time spent with four-limb support (grey boxes) in Hes2 neuron-silenced mice compared to controls. HR=hindlimb right, FR=forelimb right, HL=hindlimb left, FL=forelimb left. Scale bar is 100 msec. (C) Bar graph showing the significant increase in the time spent in four-paw support in Hes2 neuron-silenced mice compared to controls. (D,E) Bar graphs showing the significant increase in the latency of initiation of swing between diagonal (D) and ipsilateral (E) limbs in Hes2 neuron-silenced mice compared to controls. (F,G) Representative frames showing the hindpaw trajectory during swing in control (F) and Hes2 neuron-silenced (G) mice during self-paced walking on a wide runaway. (H,I) Line graphs showing the knee, ankle, and paw kinematics during a normalized step-cycle of control (F) and Hes2 neuron-silenced (G) mice during self-paced walking on a wide runaway. SEM is indicated as shaded lines, and calculated from multiple mice (Control N=4, Silenced N=5). Statistical analysis was done using two-way ANOVA, followed by Tukey’s post hoc test. (J,K) Bar graphs showing the increased number of slips in Hes2 neuron-silenced mice compared to controls as mice cross an uneven ladder (J) or a narrow beam (K). Data are presented as mean ± SEM. Each mouse analyzed is represented with a gray filled circle. Statistical analysis, unless otherwise indicated, was done using two-tailed Student’s *t-test*.

We then analyzed how the dynamics of swing initiation across leg pairs are modified to enable these four-paw support phases in mice with Hes2 neuron inactivation. Bilateral and ipsilateral limbs alternate their swing initiation, but swing initiates simultaneously in diagonal limbs **(Figure 5B)**. In Hes2 neuron-silenced mice we observed no changes in the timing of swing initiation between bilateral (**Figure S4E** and **S4F**) or diagonal limbs **(Figure 5D)**, but we saw a delay in the initiation of swing between ipsilateral limbs **(Figure 5E)**. Consistent with these observations, we recorded an altered temporal pattern of contraction of ipsilateral forelimb and hindlimb muscles (**Figure S4G-S4I**). In control mice the bursting of the tibialis anterior (hindlimb flexor, marking initiation of swing) is followed by the bursting of the ipsilateral triceps brachii (forelimb extensor, marking initiation of stance). In contrast, Hes2 neuron-silenced mice have a longer latency between the contraction of the tibialis and the triceps muscles (**Figure S4G-S4I**). Taken together, these results suggest that the increased ipsilateral latency may be the driver for the increased time mice spend in the four-paw support phase.

Next, we investigated how these new ipsilateral spatiotemporal dynamics affect the kinematics of hindlimb movement following the silencing of Hes2 neurons. We tracked the position of foot, ankle and knee in Hes2 neuron-silenced mice compared to littermate controls (**Figure 5F–5I**). After normalizing the step cycle length, we observed that inactivation of Hes2 neurons caused a significant shift in the foot and ankle trajectories during the swing and stance phase, respectively. Controls propelled the paw forward (targeting phase) mid-swing until the foot approached the forelimb position, whereas Hes2 neuron-silenced mice rapidly placed the paw back to the ground, failing to execute the targeting phase and prematurely terminating the swing phase (**Figure 5F–5I**). This premature termination also determined an earlier upward movement of the ankle at the beginning of stance (**Figure 5H–5I**).

Taken together, our observations show that silencing Hes2 neurons is sufficient to modify the temporal and spatial dynamics of ipsilateral forelimb and hindlimb coordination. Mice adopt a new way of walking that generates compensatory four-paw support intervals and suppresses the balance-challenging muscle synergies, such as the hindlimb targeting phase during mid-swing.

### Hes2 neurons ensure the correct execution of challenging motor tasks

We reasoned that the impaired ability of Hes2 neuron-silenced mice to coordinate ipsilateral limbs should alter their performance during more complex motor tasks that require balance and precise paw placement. This was assessed by setting-up three behavioral tasks, in which mice have to navigate increasingly challenging tasks and rapidly correct the ongoing movements to prevent slips and maintain balance (Gatto et al., 2021). First, we let mice, which were not previously trained on the task, cross an uneven ladder and measured the number of slips of their forelimbs or hindlimbs. While control mice navigated the asymmetric ladder by correctly placing their paws on the rungs, Hes2 neuron-silenced mice displayed an increased number of slips **(Figure 5J)**.

Next, we tested the coordination of mice in an additional task that requires both balance and precision of foot placement, namely crossing a rectangular narrow beam. Hes2 neuron-silenced animals navigated the larger beam with almost no errors, similar to controls. However, when challenged with a narrower beam, they displayed an increased number of slips while performing the task **(Figure 5K)**. Finally, mice were challenged with a circular narrow beam, which provided an additional level of difficulty due to the inability of mice to grasp the edge of the beam for better stability. Once again, Hes2 neuron-silenced mice displayed an even higher number of slips (**Figure S4J**), and were less adept at correcting their gait to smoothly perform this task.

Taken together, these findings show that the Hes2 neurons, which include the spinal V2 neuron population, are necessary for ensuring the coordination needed to perform challenging motor tasks that require the rapid integration of sensory feedback to appropriately adjust body coordination.

### Spinal V2 neurons coordinate forelimb and hindlimb movements

Interpretation of the Cre-dependent silencing of Hes2 neurons can be confounded by (1) developmental compensation and (2) the potential involvement of the brainstem and hindbrain neurons targeted by *Hes2^iCre^*. To directly address the role of V2 neurons within the adult spinal circuits, we undertook an intersectional genetic approach by crossing the *Hes2^iCre^*with the *hCdx2::FlpO* line (Britz et al., 2015) and the Cre- and FlpO-dependent DTR effector line, *Tau^ds-DTR^* (Duan et al., 2014) **(Figure 6A)**. This intersectional approach restricts V2 neuron ablation to the spinal cord following injection of the Diphteria Toxin (DT) in adult mice (P28), also ensuring the animals developed with intact spinal circuits **(Figure 6A)**. As control, we used littermates carrying all the alleles but injected with saline.

**Figure 6.**
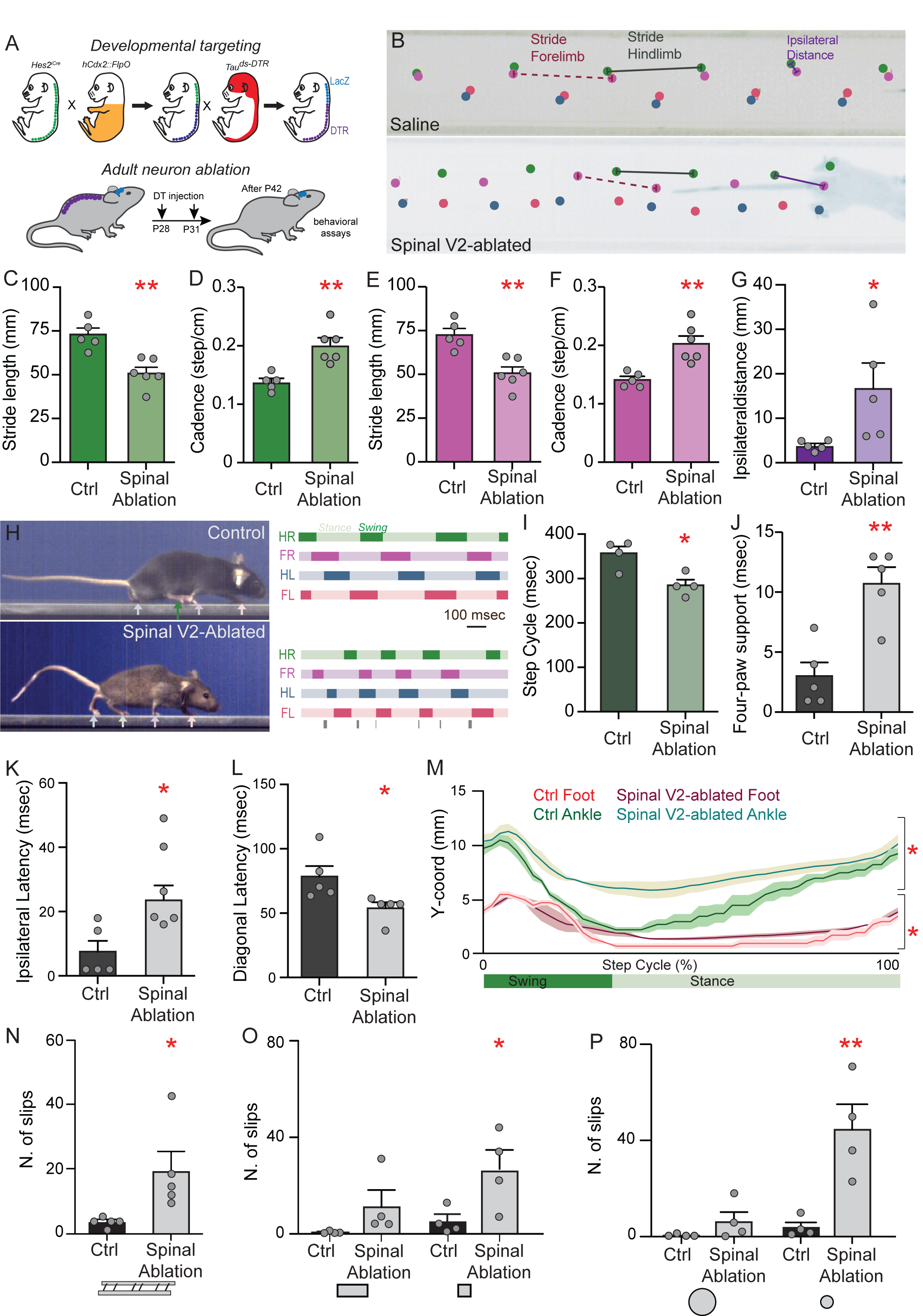
Ablation of spinal V2 neurons in adult induces festination and impairs ipsilateral body coordination. (A) Schematic illustrating the intersectional genetic approach to ablate the spinal V2 neurons in adult mice. (B) Bottom view of mice during self-paced walking on a runway. Paws are tracked in the indicated colors in control (upper panel) and spinal V2 neuron-ablated (bottom panel) mice. Stride and ipsilateral distance are calculated as indicated. (C,E) Bar graphs showing the significant shortening of the stride length of hindlimb (C) and forelimb (E) in spinal V2 neuron-ablated mice compared to controls. (D,F) Bar graphs showing the significant increased cadence of hindlimb (D) and forelimb (F) stepping in spinal V2 neuron-ablated mice compared to controls. (G) Bar graph showing the increased ipsilateral distance in spinal V2 neuron-ablated mice compared to controls. (H) Representative frames showing limb positioning in control (upper panel) and spinal V2 neuron-ablated (bottom panel) mice during self-paced walking on a wide runaway. Dark and light green arrows point to the hindlimb in stance and swing position, respectively. Representative schematic showing the temporal dynamics of interlimb coordination during self-paced walking. Note the extended time spent with four-limb support (grey boxes) in V2 neuron-ablated mice compared to controls. HR=hindlimb right, FR=forelimb right, HL=hindlimb left, FL=forelimb left. Scale bar is 100 msec. (I) Bar graph showing the significant shortening of the step cycle duration of the hindlimb in spinal V2 neuron-ablated mice compared to controls. (J) Bar graph showing the significant increase in the time spent on four-paw support in spinal V2 neuron-ablated mice compared to controls. (K,L) Bar graphs showing the significant changes in the latency of initiation of swing between ipsilateral (K) and diagonal (L) limbs in spinal V2 neuron-ablated mice compared to controls. (M) Line graphs showing the ankle and paw kinematics during a normalized step-cycle of control and spinal V2 neuron-ablated mice during self-paced walking on a wide runaway. SEM is indicated as shaded lines, and calculated from multiple mice (Control N=4, Silenced N=3). Statistical analysis was done using two-way ANOVA, followed by Tukey’s post hoc test. (N-P) Bar graphs showing the increased number of slips in V2 neuron-ablated mice compared to controls as mice cross an uneven ladder (N), a narrow beam (O) and a circular narrow beam (P). Data are presented as mean ± SEM. Each mouse analyzed is represented with a gray filled circle. Statistical analysis was done, unless otherwise indicated, using two-tailed Student’s *t-test*.

Importantly, the selective ablation of the spinal V2 neurons in adult mice recapitulated the festination phenotype observed in Hes2 neuron-silenced mice **(Figure 6B-F, Videos S3** and **S4**). Specifically, ablating the spinal V2 neurons shortened the stride and increased the cadence of both hindlimb and forelimb (**Figure 6B–6F**). Sway length, contrary to what observed in mice with developmentally silenced Hes2 neurons, was not altered following spinal V2 neuron ablation in adult animals (**Figure S5B**). Nevertheless, as previously observed for the Hes2 neuron silencing, ablation of the spinal V2 neurons caused an increase in the ipsilateral limb distance **(Figure 6G)** and a decreased distance between diagonal limbs (**Figure S5C**). Thus, the inactivation of the spinal V2 neurons is sufficient to phenocopy the gait phenotype observed in Hes2 neuron-silenced mice. Next, we analyzed how spinal V2 neuron ablation affects intra- and interlimb temporal dynamics **(Figure 6H)**. Spinal V2 neuron ablation recapitulated the hindlimb phenotypes, with reduced duration of the step cycle, swing and stance but normal stance/swing ratio **(Figure 6I, S5D-S5F**), however in forelimbs only caused a shortening of the swing phase (**Figure S5G-S5I**). Consistent with the developmental silencing results, spinal V2 neuron-ablated mice walked with more intervals of four-paw support (**Figure 6H** and **6J**), similarly due to changes in the temporal dynamics of ipsilateral limb coupling **(Figure 6K)**. Contrary to what observed in Hes2 neuron-silenced mice, following spinal V2 ablation, forelimbs are only partially affected, thus causing an additional aberrant bilateral and diagonal limb phasing **(Figure 6L** and **S5J-K**).

We then assessed whether the ablation of spinal V2 neurons phenocopied the lack of hindlimb targeting phase as observed following Hes2 neuron-silencing. We reconstructed the kinematics of foot and ankle movements during a normalized step cycle, observing that spinal V2-ablated mice prematurely terminated hindlimb swing without executing the targeting phase (**Figure 6M** and **S5L-M**). Taken together our data suggest that spinal V2 neurons are needed for the hindlimb targeting phase occurring mid-swing.

Finally, we also confirmed that ablation of the spinal V2 neurons was sufficient to impair the smooth execution of challenging motor tasks, as ablated mice displayed increased numbers of slips when crossing the asymmetric ladder **(Figure 6N)**, and the narrow **(Figure 6O)** and circular narrow beams **(Figure 6P)**.

Taken together, these data show that an intact spinal V2 microcircuit is essential for normal ipsilateral forelimb and hindlimb coordination. Moreover, these findings demonstrate that the gait alterations observed following the developmental silencing of Hes2 neurons are caused by the malfunction of spinal V2 neurons, and are unlikely due to adaptation mechanisms or inactivation of Hes2 neurons in supraspinal centers. Finally, our intersectional genetic approach demonstrates that spinal excitatory V2a and inhibitory V2b neurons function synergistically to secure ipsilateral interlimb coordination and body stability during locomotion.

## Discussion

This study reveals how excitatory and inhibitory neurons derived from a common spinal progenitor domain, the V2 lineage, cooperate to regulate the diverse motor coordination layers underlying locomotion. Our in-depth analyses of intra-and interlimb coordination demonstrate that excitatory V2a and inhibitory V2b neurons display a dual-mode of operation by exerting opposing actions in controlling flexor/extensor alternation but functioning synergistically to coordinate ipsilateral body movements. Simultaneous disruption of spinal V2a and V2b neuron activity impairs the coordination of forelimb and hindlimb muscles resulting in body instability that, although compensated during self-paced locomotion by the adoption of a festinating gait, hinders the performance in skilled locomotor tasks, such as crossing a narrow beam.

### Contribution of supraspinal vs spinal circuits to ipsilateral body coordination

In restricting Hes2 inactivation to the spinal cord, we were able to phenocopy the major aspects of motor impairment observed in Hes2 neuron-silenced mice, namely the festination-like gait, the increased four-paw support, the lack of hindlimb forward targeting, and the inability to execute skilled locomotor tasks. Some motor deficits, however, were not fully recapitulated by our spinal ablation, suggesting a) the functional contribution of supraspinal neuron populations targeted by *Hes2^iCre^*, b) compensatory changes in the spinal circuitry following Hes2-neuron developmental silencing, or c) incomplete intersectional recombination at upper cervical levels, which results in residual V2a and V2b neuron functionality. Previously we found that the *hCdx2::FlpO* allele targets the caudal spinal cord, with recombination of forelimb motor circuits occurring with a salt and pepper pattern between C2 and C3 segments (Britz et al., 2015). This is consistent with spinal V2-neuron ablation, compared to Hes2 neuron silencing, causing a less severe motor deficit in forelimbs (**Figure S5G-S5I**), affecting only the swing duration, (**Figure S3G-S3I**), but fully recapitulating the hindlimb phenotype **(Figure 6I, S5D-S5F)**. The reduced base of support (hindlimb sway) in Hes2-neuron silenced mice (**Figure S4B**) but not in spinal V2-neuron ablated mice (**Figure S5B**) is also suggestive of residual functionality at cervical level, as reported in spinal contusion models, where reductions in hindlimb sway are scaled to the severity of cervical damage (Beare et al., 2009).

The contribution, if any, to interlimb coordination of the ventral PAG, the pons, and the superior olive captured by *Hes2^iCre^*(**Figure S2G**) is not well understood. However, we speculate that *Hes2^iCre^*targeting of neurons in the medullary reticular nucleus, which contains spinally projecting V2a neurons (Bouvier et al., 2015; Cregg et al., 2020; Usseglio et al., 2020), might also account for the partial deficits in the forelimbs of spinal V2 neuron-ablated mice.

In summary, our results identify a key spinal microcircuit that synchronizes ipsilateral body movements, thereby confirming prior lesion studies ascribing the control of interlimb coordination to the spinal circuitry (Miller et al., 1975), with supraspinal circuits exerting phase-dependent modulatory effects (Rossignol et al., 1993).

### Distinct cooperation dynamics of excitatory V2a and inhibitory V2b neurons underlie different facets of limb coordination

Previous studies suggest the net balance of excitatory and inhibitory input dictates the firing pattern of motoneurons, and the consequent contraction of muscles (Petersen et al., 2014; Ramírez-Jarquín and Tapia, 2018). Nevertheless, it remains largely unclear how this input is distributed across motoneuron types, e.g. flexor and extensor, and how is coordinated within and across spinal segments. Spinal half-center modules are organized in reciprocally coupled flexor and extensor units, with V1 and V2b neurons preferentially innervating flexor and extensor motoneurons, respectively (Britz et al., 2015). While inactivation of V2b neurons induces hyper-extension of the hindlimb (Britz et al., 2015), inactivating excitatory V2a neurons does not alter flexor/extensor patterning (Crone et al., 2009). Interestingly, the concomitant inactivation of excitatory V2a and inhibitory V2b neurons rescues the hyper-extension phenotype (**Figure 5F–5I, S3J-S3L**) observed in V2b neuron-ablated mice (Britz et al., 2015), suggesting that these populations act with opposing effects on the recruitment of extensor muscles. Clonally related V2a and V2b neurons predominantly synapse onto distinct target neurons in zebrafish, providing a cellular logic for the segregated circuit integrations of V2a and V2b neurons (Bello-Rojas and Bagnall, 2022).

The coordination of hindlimb and forelimb seems to involve a different cooperation dynamic compared to the flexor/extensor alternation, whereby the simultaneous inactivation of V2a and V2b neurons introduces a new motor deficit, the aberrant ipsilateral body coordination (**Figures 5** and **6**), not observed when these populations are individually ablated (Britz et al., 2015; Crone et al., 2009). This raises the question of how this synergistic interaction is encoded at a circuit level. The forelimb and hindlimb CPGs for walking are connected by propriospinal ascending and descending neurons (Grillner and El Manira, 2020). These propriospinal pathways include excitatory and inhibitory neurons, which elicit both EPSP and IPSP in the post-synaptic premotor neurons or motoneurons (Jankowska et al., 1973, 1974; Lloyd and McIntrye, 1948). As both V2a and V2b comprise subsets of propriospinal neurons (Flynn et al., 2017; Ruder et al., 2016), it is intriguing to speculate that these neurons regulate interlimb coordination by acting as stop and go signals to harmonize the initiation and termination of motoneuron firing across segments.

The differential effects that the simultaneously inactivation of V2a and V2b neurons has on flexor/extensor (intralimb) *vs* forelimb/hindlimb (ipsilateral) coordination suggest a unique duality of synergistic and balanced interactions. An intriguing hypothesis to explain this duality of function is that specialized subsets within the V2a and V2b neuronal populations underlie local vs segmental motoneuron coordination. V2a and V2b neurons comprise two subtypes labeled by Nfib, NeuroD2, and Prox1 (N-type) and Zfhx3, Zfhx4, and FoxP2 (Z-type), which mark local vs projection neurons, respectively (Osseward et al., 2021). Functionally targeting these subtypes will be key to untangle the circuits underlying flexor/extensor and ipsilateral limb coordination, as well as their distinct dynamics of operation.

### Circuits for limb coordination and postural stability

The emergence of a festination-like gait is the most prominent deficit that occurs upon inactivating or ablating Hes2 neurons in the spinal cord. Festination is characterized by a combined increase in the cadence of stepping and reduction in stride length (**Figures 4A–4F** and **6B-6F**), and it is remarkably similar to the abnormal gait adopted in Parkinsońs patients. Strikingly, the deficits in hindlimb targeting observed in mice with spinal V2 neuron inactivation mirror the reduced forward limb propulsion reported in Parkinsońs patients (Knutsson, 1972; Morris et al., 1999) (**Figures 5F–5I, 6M** and **S5L-S5M**). Parkinson’s patients typically increase the double support phase during stepping (Arippa et al., 2022), which is also comparable to the prolonged four-paw support phase we observe following spinal V2 neuron silencing or ablation (**Figures 5B–5C** and **6H,6J**). Furthermore, the absence of forelimb and hindlimb coordination in V2 neuron-inactivated mice (**Figures 5** and **6**) mimics the reduction in arm swinging observed in Parkinson’s patients, especially during challenging motor tasks (Baron et al., 2018; Siragy et al., 2021). Finally, Parkinsońs patients are at high risk of falling or losing balance, especially during turning or dual-tasking (Mirelman et al., 2019), which is remarkably similar to the increase in foot slips observed in V2 neuron-inactivated mice during skilled locomotor tasks (**Figure 5J, 5K, S4J** and **6N-6P**).

While the primary cause of the movement deficit in Parkinson’s patients is closely tied to the degeneration of the basal ganglia circuitry, the Hes2 neuron inactivation-induced festination arises from disrupting the spinal cord circuitry. Deficits in dopamine signaling in the basal ganglia may have a direct effect on V2 neurons, which lie downstream of the cortico-basal ganglia network (Leiras et al., 2022). Alternatively, the festination motor phenotype might arise as a compensatory mechanism to maintain balance while walking, with the common motif across Parkinson and V2 neuron inactivation being the emergence of postural instability. Interestingly, Parkinson’s patients develop an abnormal forward flexion of the trunk, which destabilizes their center of mass (Morris et al., 1999). The underlying nature of postural instability in V2 neuron-ablated mice is less obvious, in part, because they are quadrupeds. An intriguing possibility is that propriospinal pathways coordinate motor activity across multiple spinal segments to stabilize body balance, and inactivation of these pathways leads to an unstable posture. Propriospinal pathway are critical to relay sensory and supraspinal feedback across segments (Alstermark et al., 1987, 2007; Brockett et al., 2013; Ruder et al., 2016), making it appealing to speculate that these neurons sense limb stability, and following integration of mechanoreceptive and proprioceptive feedback, direct the ipsilateral limb to initiate/end the swing phase. Consistently, in mice and insects it has been shown that the posterior limb is guided forward by the position of the anterior ipsilateral limb via proprioceptive feedback (Brunn and Dean, 1994; Mayer and Akay, 2021; Theunissen et al., 2014). V2a and V2b neurons receive extensive sensory input from proprioceptive and mechanoreceptive afferents (Di Costanzo, Stam, Goulding unpublished data), and therefore the way they integrate and relay these sensory inputs might be at the core of their contribution to postural stability.

In summary, our findings show that the collective inactivation of spinal V2 neurons destabilizes ipsilateral movements and leads to similar compensatory mechanisms as what observed in Parkinson’s patients. These similarities might be due to V2 neurons acting downstream of the basal ganglia or by the occurrence of similar postural unbalance in these two conditions. Parkinsońs patients display an array of motor impairments (Mirelman et al., 2019), responsive to a different degree to the treatment with dopamine agonist. While freezing of gait can be ameliorated by levodopa treatment, the increased cadence and shortened stride, although present from the initial stages of the disease, are often non-responsive to this treatment (Galna et al., 2015). Thus, spinal V2 neurons could be a new therapeutic target, as their pharmacological or electrical stimulation might widen the spectrum of treatable motor symptoms, by providing alternative strategies to replace the lost descending inputs or to steady the body posture.

## Materials and Methods

### Mice

Mice were maintained following the protocols for animal experiments approved by the IACUC of the Salk Institute for Biological Studies according to NIH guidelines for animal experimentation and of the Kyoto University (permit numbers: Med Kyo 16216 and Lif-K18018). 6-to 20-week old mice of both sexes were used for behavioral experiments. Analysis of the behavioral data showed no gender-bias, with similar responses observed in male and female mice. E9.5 to 8-week old mice (as indicated in the respective method paragraphs) were used for anatomical and molecular characterization of the spinal neurons. Mice were housed in groups of up to 5 animals on a 12-hour dark/light cycle in a humidity-and temperature-controlled room, and provided with rodent diet (Picolab) and water ad libitum. The following strains of mice were used: *R26^Ai14-LSL-tdTomato^*(Madisen et al., 2010) (JAX, 007908), *R26^LSL-Sun1::EGFP^* (JAX, 201039), *R26^LSL-LacZ^* (Soriano, 1999), *R26^LSL-TenT^* (Zhang et al., 2008), *Tau^ds-DTR^* (Duan et al., 2014), *Gata3*^nlsLacZ^ (van Doorninck et al., 1999), *hCdx2::FlpO* (Bourane et al., 2015), *R26^Ai65-LSL-FSF-tdTomato^* (Madisen et al., 2015).

### Generation of the Hes2^iCre^ mouse line

To construct the *Hes2-iCre* knock-in targeting vector, the *iCre* coding sequence, the SV40-derived polyadenylation sequence (pA), and the FRT-flanked neomycin selection cassette were inserted into the translation initiation site of *Hes2* (Isaka et al., 1996). This engineered fragment was inserted into the pBluescript II SK+ vector (Stratagene) with a DTA-negative selection cassette and Hes2 5’ and 3’ homology arms of 3.2-kb and 6.2-kb, respectively. The targeting vector was linearized with NotI and electroporated into mouse TT2 embryonic stem cells, and G-418 resistant clones were selected. Genomic DNA from drug-resistant cells was digested with XmnI and analyzed by Southern blot using a 0.3-kb DNA fragment as a 5’ external probe for Hes2 (**Figure S2A**). Chimeric mice were produced from two successfully targeted ES cell clones by aggregation with ICR embryos. Germ line transmission of the targeted allele was assessed by PCR of tail DNA. Neomycin selection cassette was removed by crossing with *pCAG-FLPe* mice (Kanki et al., 2006). Subsequently, a PCR strategy was used to identify the mutants. Genotypes were determined by PCR using the following primers:

Hes2-WT-fw, 5’-AGCCTAGTGGCTGATAGTGAGCG −3’;

Hes2-WT-rev, 5’-ACCACAGTTAGAAAGACCGCCATCG −3’;

Hes2-KO-rev, 5’-AGGCCAGATCTCCTGTGCAGCATG −3’.

The PCR product sizes for mutant allele and wild-type allele are 269 bp and 507 bp, respectively.

### In ovo electroporation

The CAG:Hes2-IRES-GFP plasmid was generated as follows. Hes2 coding sequence was PCR-amplified from a cDNA library generated from E12 mouse embryos. The Hes2 sequence was cloned into pCR-TOPOII vector (ThermoFisher). After confirming the sequence via Sanger sequencing, the Hes2 sequence was cloned into CAG:IRES-GFP plasmid (Addgene #11159).

Chick eggs (Charles River and McIntyre Farms) were incubated in a humidified chamber. Hes2 expression construct (1 ug/μl in PBS and fast green) was injected into the lumen of Hamburger and Hamilton (HH) stage 12-14 chick developing neural tube (Hamburger and Hamilton, 1951). Electroporation was performed using a square wave electroporator (BTX). Incubated chick embryos were harvested after 48 hr.

### Whole Mount Staining

Dissected E10.5 and E11.5 embryos were fixed in 2% formaldehyde, 0.2% glutaraldehyde in PBS solution for 60 min at 4°C, rinsed twice in 0.1 M phosphate buffer (pH 7.4), 2mM MgCl2, 0.01% sodium deoxycholate, 0.02% NP-40. Embryos were then stained at 37°C for 8 hr in X-gal stain buffer (2 mg X-gal in 2mM MgCl2, 0.01% sodium deoxycholate, 0.02% NP-40, 5mM K3Fe(CN)6, 5mM K4Fe(CN)6 in 0.1M phosphate buffer), as described previously (Imayoshi et al., 2006). Stained embryos were washed twice in PBS and post-fixed with 4% PFA in PBS for 2 hr at room temperature.

### Immunohistochemistry

Embryos were fixed with 2-4% PFA for 60-120 min at 4°C. Postnatal spinal cords were isolated and fixed with 4% PFA for 60-120 min at 4°C. Tissues were washed with PBS, equilibrated in 30% sucrose for 2 hr at 4°C, embedded in Tissue-Tek OCT (Sakura), and subjected for cryosectioning onto glass slides (VWR). Immunohistochemistry was performed by incubating with primary antibodies (overnight, 4°C) and fluorophore-conjugated secondary antibodies (2 hr, room temperature; ThermoFisher, Jackson ImmunoResearch). Sections were mounted with VectaShield (VECTOR) or Mowiol and coverslipped (VWR). Images were acquired using an Olympus FV1000 and FV3000. Images are presented as z-projections unless otherwise noted.

The following primary antibodies were used: guinea pig anti-Chx10 (1:2000) (Thaler et al., 2002), rabbit anti-βgal (1: 5000; Cappel), rabbit anti-HB9 (1:6000) (Thaler et al., 1999), goat anti-GFP (1:1000; Rockland). The following secondary antibodies were used: Donkey anti-rabbit (ThermoFisher, 1:1000), anti-goat (ThermoFisher, 1:1000), and anti-guinea pig (Jackson Immuno, 1:300).

### In situ hybridization

Head and tail of embryos were removed, and the remaining tissue was fixed in ice-cold 4% PFA for 1 hr at 4 °C. After a brief PBS wash and 2 hr of 30 % sucrose incubation at 4 °C, embryos were cut in half through the mid-thoracic cord level, embedded in OCT, frozen, and stored in −80 °C. Following cryosectioning (16 μm), RNAscope Fluorescent Multiplex Assay v1 was performed according to the manufacturer’s instructions, except 1:2 dilution of protease IV in PBS was applied for 20 min. Following the RNAscope, sections were subjected to immunohistochemistry, with blocking buffer (1% BSA in 0.1M Triton-X/PBS) for 1 hr, primary antibodies (rabbit anti-Hb9 (1:8000) and guinea pig anti-Chx10 (1:500)) in blocking buffer overnight at 4 °C; and secondary antibodies goat anti-rabbit 488 (1:1000) and goat anti-guinea pig 647 (1:500) in blocking buffer for 2 hr at room temperature. Following DAPI treatment, Moviol was used to mount coverslips on the slides. Images were acquired with an Olympus 3000 confocal microscope.

### Neuronal ablation

Mice carrying the *Hes2^iCre^;hCdx2::FlpO* alleles in addition to the effector *Tau^ds-DTR^* received i.p. injections of diphtheria toxin (DTX, 50 ng/gram of weight; List Biological Laboratories) at P28 and P31, or saline solution in equal volume as control.

### Behavioral assays

#### Runway foot print analysis

Bottom-view videos were recorded through a transparent platform. For each mouse we acquired two trials, and one of the two videos was selected for analysis. For each mouse a minimum of five consecutive steps in the middle of the walk-through were analyzed for all the parameters measured. We registered the position of the center of each paw for each step using the ImageJ cell counter plug-in.

#### Kinematic reconstructions

The trials of mice walking on the 25 mm wide beam (see below) were used for the marker-less reconstructions of joint kinematics. Step cycle phases for each limb and the 2D joint kinematics were tracked semi-automatically using the Simi Motion Analysis System. At least four mice per genotype were used for kinematic analysis, and for each mouse at least 5 consecutive steps were analyzed.

#### Narrow Beam Test

Mice were trained on the first day to cross 3-5 times an elevated one meter long and 25 mm wide beam. No differences in performance were observed during this training. In the following days, mice were tested on the elevated narrower beams (12 mm, 5 mm rectangular beams and 25 mm and 5 mm circular beams). Individual trials were video-recorded using two high-speed cameras (mV Blue Cougar XD; 200 frame/second), capturing the performance from a lateral and a bottom view. Each mouse crossed three times each beam, and the total number of slips from all limbs in the three runs was calculated by analyzing the videos frame by frame using the Simi Motion Analysis System.

#### Ladder Test

One day after the narrow beam tests were performed, animals were tested onto the uneven ladder. Individual trials were video-recorded using two high-speed cameras (mV Blue Cougar XD; 200 frame/second), capturing the performance from a lateral and a bottom view. Each mouse crossed three times the ladder, and the starting side was alternated at each trial. The total number of slips for all the limbs in the three runs was calculated by analyzing the videos frame by frame using the Simi Motion Analysis System.

#### Grip strength

A digital grip strength meter (San Diego Instruments) was used to measure force (grams) of mouse forelimb and hindlimb grip response. For hindlimb measurements, animals were positioned to avoid contact of the forelimbs with the meter and to maximize grip reflex of the hindlimb. Five trials were performed for each animal and the 3 highest force exerted were averaged.

#### Rotarod

Animals were trained on the Rotarod on the first day for 1 minute at 3rpm. On the following three days, mice performed four trials, in which the Rotarod was accelerating from 0-50 rpm over 5 minutes. Trials were separated by 10-minute intervals. The latency to fall was recorded for each individual trial, and averages of the four trials were used to score each mouse.

#### Pinprick test

Mice were habituated on the grid for 30 minutes for two consecutive days. On testing day, they were acclimatized for 20 minutes to the grid before their hindpaws were stimulated with an Austerlitz insect pin (Tip diameter 0.02 mm; Fine Science Tools). The pin was gently applied to the plantar surface of the paw, alternating between left and right hindpaws. The stimulation was repeated 10 times, with at least one minute interval between trials. The percentage of withdrawal reflex responses elicited out of the total number of pinprick stimulations was calculated.

#### Hargreaves test

Animals were habituated in a plastic box on a glass surface for two days. Thermal pain was induced using a radiant heat beam (IITC) focused on the hindpaw of the mouse. Intensity of the heat beam was adjusted so that the withdrawal response occurred within a range of ∼3-8 seconds. Mice were never stimulated for more than 20 seconds to prevent tissue damage. Latency to exhibit a withdrawal response was measured for every trial. Two trials were performed on each hindlimb, and the average of four trials (two trials for left and right hindlimbs) was used to score each individual.

### EMG recording

EMG electrodes were fabricated as described in Pearson, Acharya, and Fouad 2005. 3-to 5-month-old mice had EMG electrode implanted in the triceps brachii and the biceps brachii of the right forelimb and the tibialis anterior and the gastrocnemius of the right hindlimb. The electrodes were stabilized in place with a knot on the distal end and the skin was sutured to close the wound. The headpiece was cemented on the skull of the mouse, and then attached to an adaptot to transmit the signal to the pre-amplifiers (MA103, University of Cologne). The signal was then filtered (low pass 10 Hz – high pass 10 kHz) by the amplifier (MA102, University of Cologne), and digitized with a Micro1401 Digitizer (CED). The EMG signals were recorded and analyzed using the Spike2 software. The raw signals were rectified and integrated, and latency was determined based on the first spike.

## Acknowledgements

We thank members of the NIH U19 - Spinal Circuits for the Control of Dexterous Movement for helpful discussions on experiments and analyses. This work was funded by the National Institutes of Health (NIH-U19NS112959 to M.G. and S.L.P.) and the German Research Foundation (DFG CRC1451 to G.G.).

## Supplemental Figure Legend

**Figure S1. Identification of a novel marker of the spinal V2 neuron lineage**

(A) A t-distributed stochastic neighbor embedding plot representing neuron clusters in the developing spinal cord (Delile et al., 2019) and showing the expression of *Lhx3* and *Hes2*. Red circles highlight the cluster that co-expresses *Lhx3* and *Hes2*. Data mined (tSNE Plot, Perplexity=20, K=33) via the Single Cell Expression Atlas website <u>Single Cell Expression Atlas −</u> <u>EMBL-EBI</u>

(B,C) Cross-sections of the E10.5 spinal cord showing the segregated expression of *Hes2* (violet) and motoneuron-specific probes *Isl1* (red) (B) or *Olig2* (red) (C). Scale bar: 100 μm.

**Figure S2. Establishing genetic access to the Hes2 neurons**

(A) Schematic showing the targeting strategy to generate the *Hes2^iCre^* knock-in line.

(B) Southern Blot to confirm the insertion of the *iCre* allele in the *Hes2* locus.

(C) Cross-sections of the brain from a postnatal day 16 *Hes2^iCre^;R26^Ai14-LSL-tdTomato^* mouse, stained with antibodies for tdTomato (red) and Neurotrace (blue). Scale bars: 500 μm.

(D) Normal ratio of wild-type, heterozygous and homozygous embryos and adult mice were found when crossing *Hes2^iCre^*heterozygous mice. Among E12.5 embryos, out of 5 litters, 13 embryos were wild-types (27%), 22 embryos were heterozygotes (47%), and 12 embryos were homozygotes (26%). Among adult mice, out of 5 litters, 9 mice were wild-types (24%), 19 mice were heterozygotes (50%), and 10 mice were homozygotes (26%).

(E,F) Bar graphs showing the absence of changes in the number of V2a (Chx10+) (E) and V2b (*Gata3^nlslacZ^*+) (F) neurons between *Hes2* heterozygous and knockout (homozygous) embryos, assessed by two-tailed Student’s t-test.

(G) Cross-sections from Hamilton Hamburger stage 23 chick embryos, which were *in ovo* electroporated at HH stage 12-14 to over-express Hes2. The sections were stained with antibodies for Gata3 (red) and Chx10 (blue). Scale bars: 100 μm.

**Figure S3. Festinating gait induced by the developmental silencing of the Hes2 neurons**

(A-D) Bar graphs showing the absence of changes between control and Hes2 neuron-silenced mice when tested on the rotarod (A,B) and for grip strength normalized by weight (C,D).

(E) Representative frames showing the alternation of stance and swing phases of the forelimb in control (upper panel) and Hes2 neuron-silenced (bottom panel) mice during self-paced walking on a wide runaway. Dark and light violet arrows point to the forelimb in stance and swing position, respectively.

(F) Representative schematic showing the reduced duration of the step cycle, stance and swing of the forelimb in Hes2 neuron-silenced mice compared to controls. Scale bar is 70 msec.

(G-I) Bar graphs showing the significant shortening of the step cycle (G), swing (H) and stance (I) duration of the forelimb in Hes2 neuron-silenced mice compared to controls.

(J) Representative EMG traces from the hindlimb flexor (Tibialis anterior – green) and the hindlimb extensor (Gastrocnemius – yellow) in control and Hes2 neuron-silenced mice.

(K) Integrated and normalized EMG signals from the hindlimb flexor (Tibialis anterior – green) and the hindlimb extensor (Gastrocnemius – yellow) in control and Hes2 neuron-silenced mice.

(L) Bar graph showing the similar latency between tibialis and gastrocnemius muscle firing in Hes2 neuron-silenced mice compared to controls.

(M,N) Bar graphs showing the absence of impairments in the execution of the withdrawal reflex induced by noxious pinprick (M) or thermal (N) stimuli in Hes2 neuron-silenced mice compared to controls.

Data are presented as mean ± SEM. Each mouse analyzed is represented with a gray filled circle. Statistical analysis was done using two-tailed Student’s *t-test*.

**Figure S4. Developmental silencing of the Hes2 neurons disrupts ipsilateral body coordination**

(A) Schematic displaying the calculation of hindlimb and forelimb sway, ipsilateral distance, and diagonal distance, with the respective color-coding for individual limb.

(B) Bar graph showing the reduced sway of hindlimb but not forelimb in Hes2 neuron-silenced mice compared to controls.

(C) Bar graph showing the reduced diagonal distance in Hes2 neuron-silenced mice compared to controls.

(D) Bar graph showing the increased ipsilateral distance in Hes2 neuron-silenced mice compared to controls.

(E,F) Bar graphs showing the reduced latency between swing initiations at hindlimb (E) but not forelimb (F) levels in Hes2 neuron-silenced mice compared to controls.

(G) Representative EMG traces from the hindlimb flexor (Tibialis anterior – green) and the forelimb extensor (Triceps Brachii – orange) in Hes2 neuron-silenced and control mice.

(H) Integrated and normalized EMG signals from the hindlimb flexor (Tibialis anterior – green) and the forelimb extensor (Triceps Brachii – orange) in Hes2 neuron-silenced and control mice. Start of hindlimb swing (green) and forelimb stance (orange) is indicated by colored rectangles.

(I) Bar graph showing the increased latency between tibialis and triceps muscle firing in Hes2 neuron-silenced mice compared to controls.

(J) Bar graphs showing the increased number of slips in Hes2 neuron-silenced mice compared to controls as mice cross a circular narrow beam.

Data are presented as mean ± SEM. Each mouse analyzed is represented with a gray filled circle. Statistical analysis was done using two-tailed Student’s *t-test*.

**Figure S5. Ablation of spinal V2 neurons induces festination and impairs ipsilateral body coordination**

(A) Schematic displaying the calculation of hindlimb and forelimb sway and diagonal distance, with the respective color-coding for each limb.

(B) Bar graphs showing no changes the sway of hindlimb and forelimb in spinal V2 neuron-ablated mice compared to controls.

(C) Bar graph showing the reduced diagonal distance in spinal V2 neuron-ablated mice compared to controls.

(D,E) Bar graphs showing the reduced swing (D) and stance (E) duration in spinal V2 neuron-ablated mice compared to controls.

(F) Bar graph showing no difference in the swing/stance ratio in spinal V2 neuron-ablated mice compared to controls.

(G-I) Bar graphs showing reduced swing (H) but no changes in step cycle (G) and stance (I) duration in forelimbs of spinal V2 neuron-ablated mice compared to controls.

(J,K) Bar graphs showing the reduced latency between swing initiations at forelimb (K) but not hindlimb (J) level in spinal V2 neuron-ablated mice compared to controls.

(L,M) Representative frames showing the hindpaw trajectory during swing in control (L) and spinal V2 neuron-ablated (M) mice during self-paced walking on a wide runaway.

