## Supplementary figures and images for "A spinal synergy of excitatory and inhibitory neurons coordinates ipsilateral body movements"

### Supplemental Figure 1

Figure S1

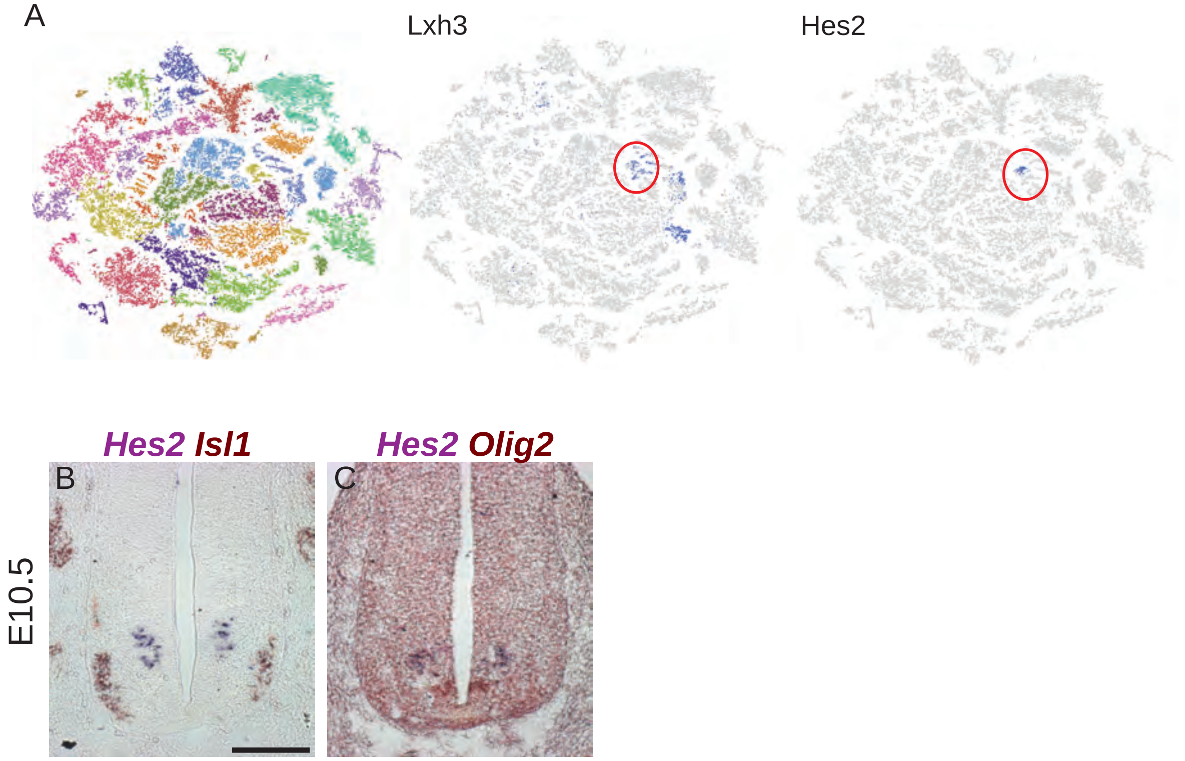

### Supplemental Figure 2

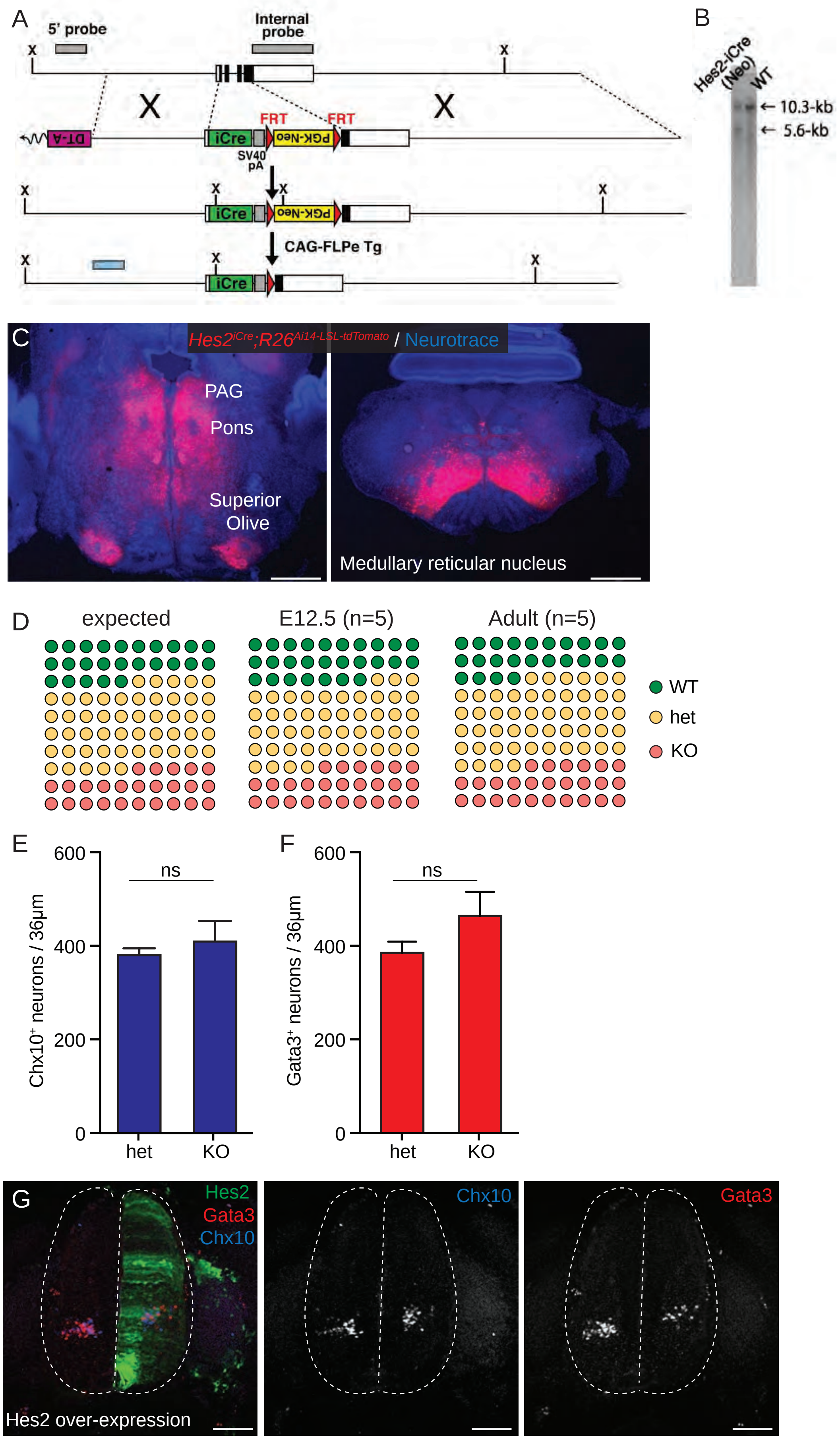

### Supplemental Figure 3

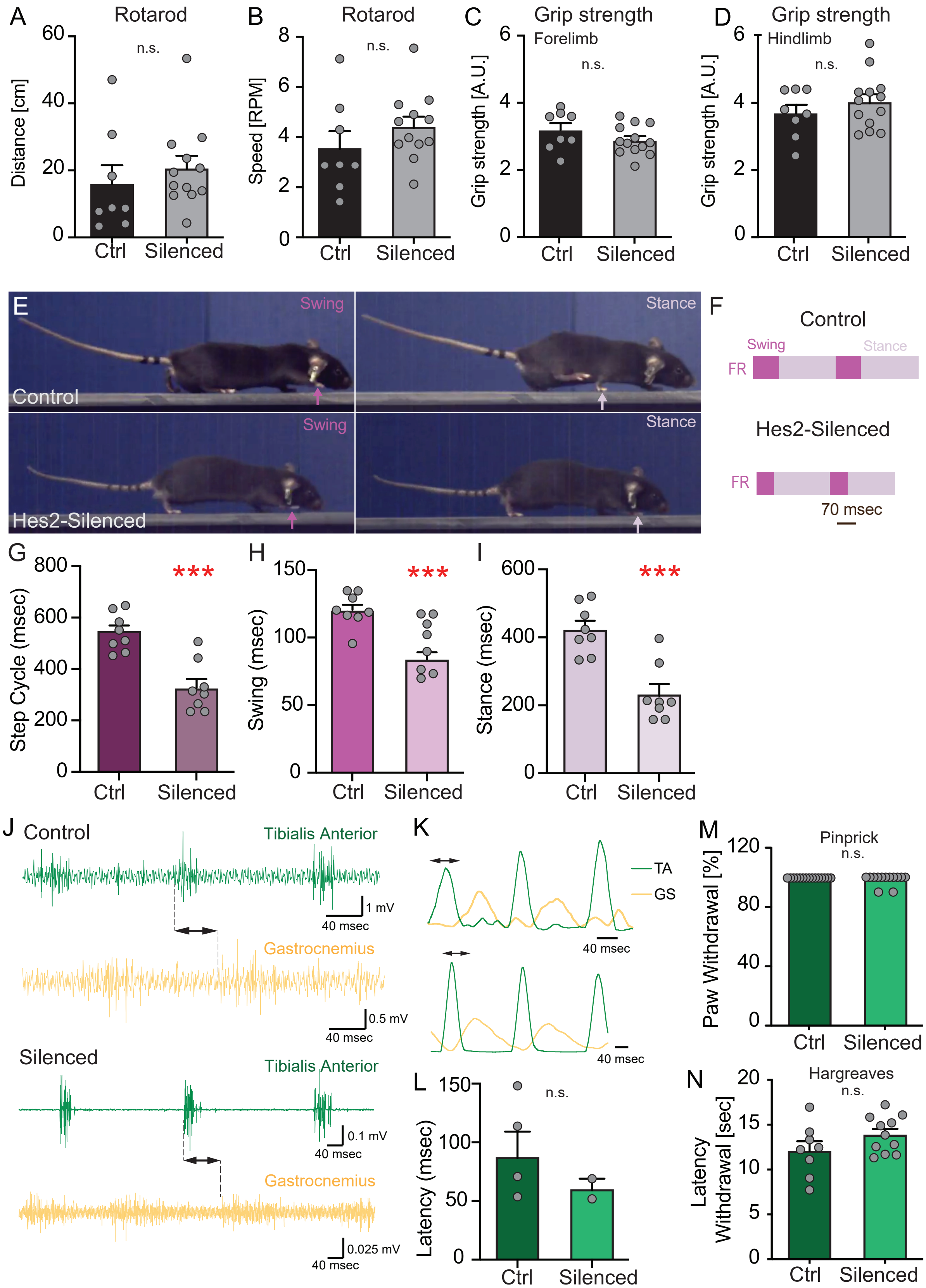

### Supplemental Figure 4

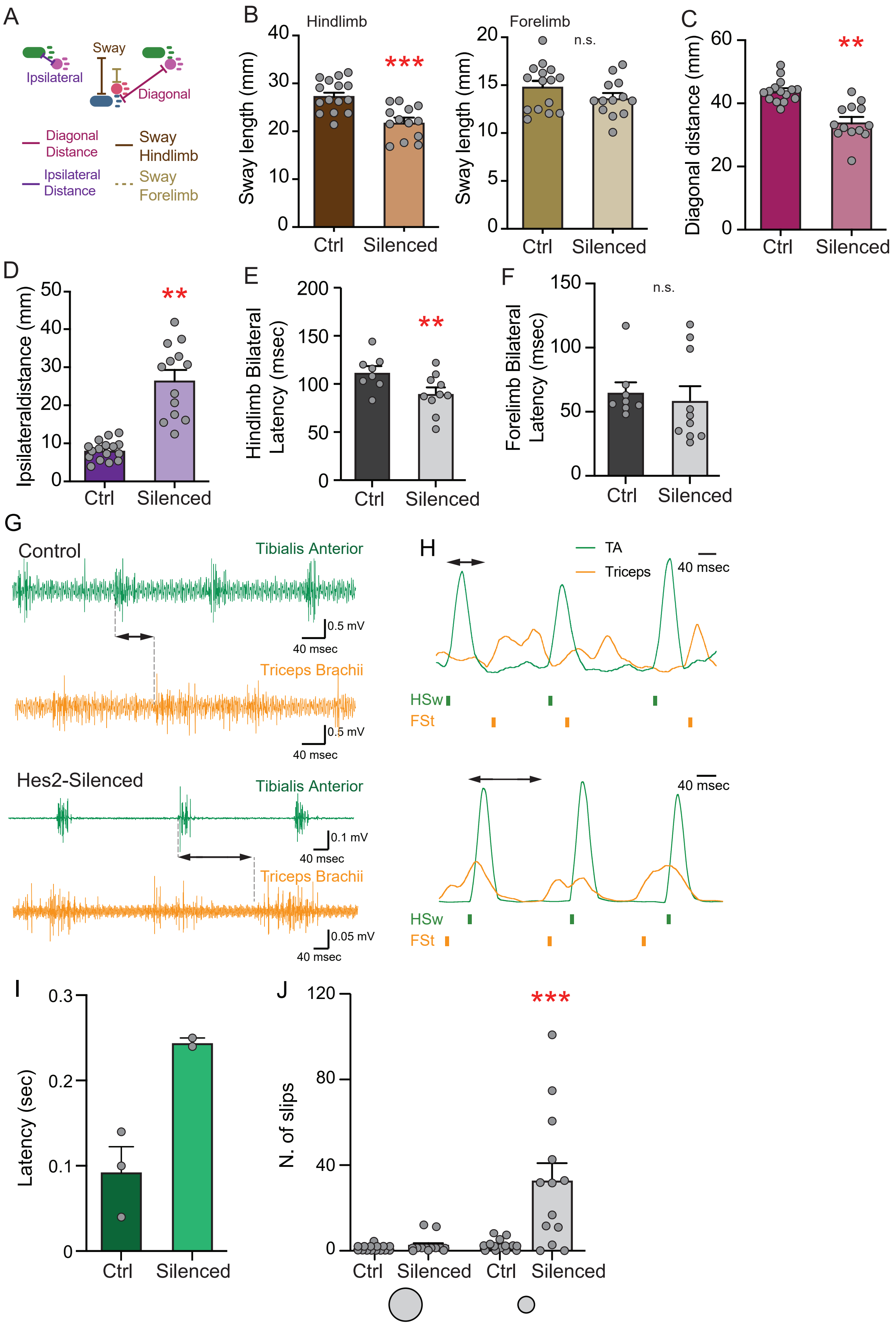

### Supplemental Figure 5

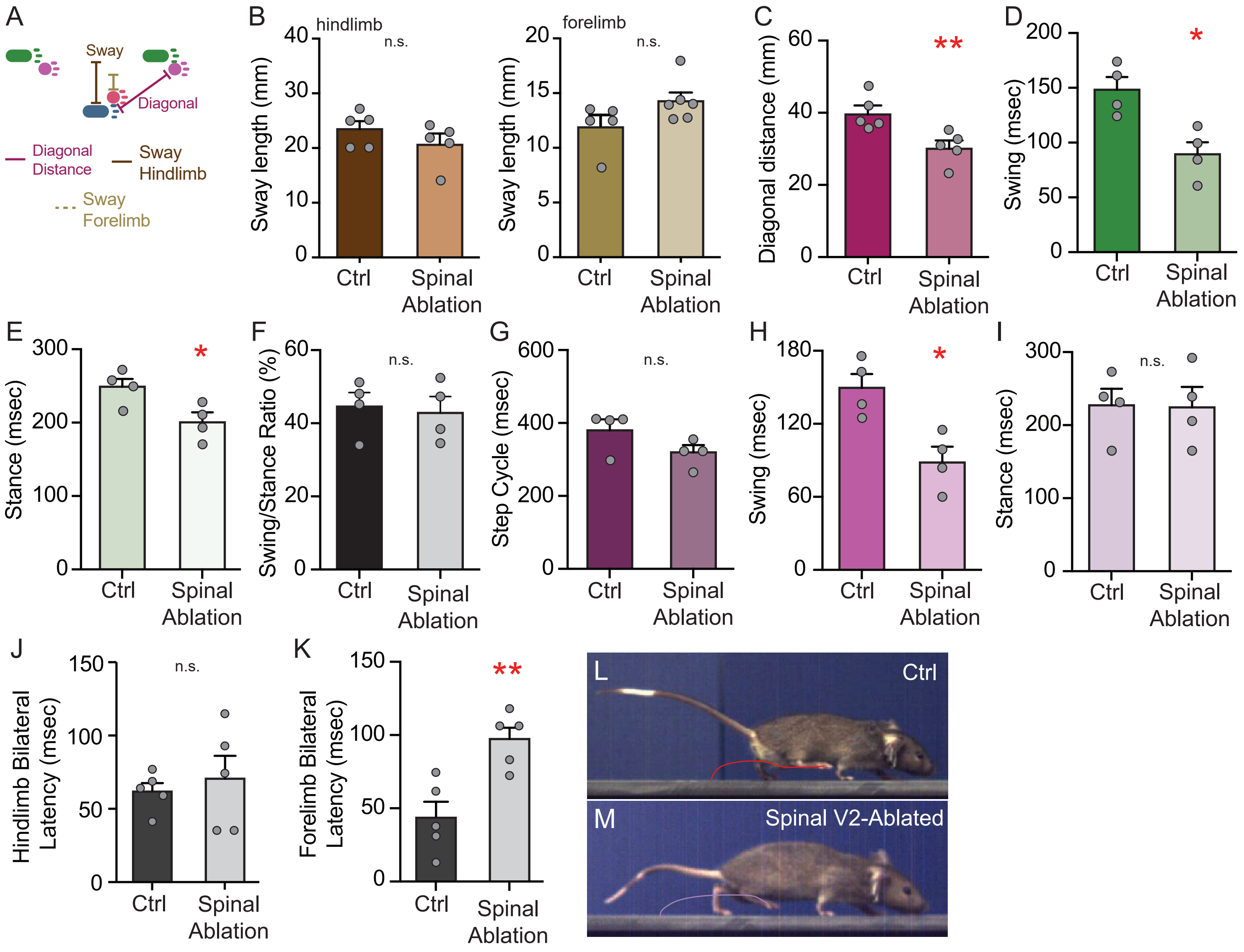
